# Population variation of miRNAs and isomiRs and their impact on human immunity to infection

**DOI:** 10.1101/2020.01.31.928580

**Authors:** Maxime Rotival, Katherine J Siddle, Martin Silvert, Julien Pothlichet, Hélène Quach, Lluis Quintana-Murci

## Abstract

MicroRNAs (miRNAs) are key epigenetic regulators of the immune system, yet their variation and contribution to intra- and inter-population differences in immune responses is poorly characterized. Here, we generated 977 miRNA-sequencing profiles from primary monocytes, from individuals of African and European ancestry, following activation of three TLR pathways (TLR4, TLR1/2 and TLR7/8) or infection with Influenza A virus. We find that immune activation leads to important modifications in the miRNA and isomiR repertoire, particularly in response to viral challenges. These changes are, however, much weaker than those observed for protein-coding genes, suggesting stronger selective constraints on the miRNA response to stimulation. This is supported by the limited genetic control of miRNA expression variability (miR-QTLs) — and the lower occurrence of G×E interactions — in stark contrast with eQTLs that are largely context-dependent. We also detect marked differences in miRNA expression between populations, which are mostly driven by non-genetic factors. Yet, on average, miR-QTLs explain ~60% of population differences in expression of their cognate miRNAs, and, in some cases, evolve adaptively, as shown in Europeans for a miRNA-rich cluster on chromosome 14. Finally, integrating miRNA and mRNA data from the same individuals, we provide evidence that the canonical model of miRNA-driven transcript degradation has a minor impact on miRNA-mRNA correlations, which are, in our setting, mainly driven by co-transcription. Together, our results shed new light onto the factors driving miRNA and isomiR diversity at the population level, and constitute a useful resource for evaluating their role in host differences of immunity to infection.

## INTRODUCTION

Since their discovery in 1993 (Lee et al. 1993), microRNAs (miRNAs) — short, evolutionary conserved RNA sequences of ~22 nucleotides — have emerged as major epigenetic regulators, involved in a large variety of developmental and cellular processes such as cell differentiation, proliferation and homeostasis (Krol et al. 2010). There is also increasing evidence that support their key role in immune responses, with miRNAs such as miR-155 or miR-146a acting to promote and stabilize the inflammatory response (O’Connell et al. 2012; Vigorito et al. 2013; Mehta and Baltimore 2016; Alivernini et al. 2017; Su et al. 2020). Furthermore, numerous studies have reported strong shifts in miRNA expression profiles in response to infectious agents, such as *Mycobacterium tuberculosis* (Siddle et al. 2014; Siddle et al. 2015), *Salmonella* (Pai et al. 2016) or Influenza A virus (Zhang et al. 2018).

Studies of miRNA abundance across various cell types and tissues have allowed the characterization of the extent of genetic regulation of miRNA expression variability, i.e., miRNA expression quantitative trait loci (miR-QTLs), and highlighted the role of genetic variants located in the promoter of the precursor transcript of the miRNA (pri-miRNAs) in shaping inter-individual differences in miRNA expression (Borel et al. 2011; Gamazon et al. 2012; Parts et al. 2012; Lappalainen et al. 2013; Siddle et al. 2014; Huan et al. 2015; Budach et al. 2016; Gottmann et al. 2018; Li et al. 2020). In the context of immunity, despite increasing evidence that marked population differences exist in the mRNA response to immune challenges (Nedelec et al. 2016; Quach et al. 2016), the extent to which miRNA responses to infection vary across individuals of different ancestry remains largely unknown.

Fueled by the advent of deep sequencing technologies, growing evidence has emerged that mature miRNAs undergo important post-transcriptional modifications (Neilsen et al. 2012; Ameres and Zamore 2013; Tan et al. 2014; Trontti et al. 2018; Kim et al. 2019). These include nucleotide substitutions (miRNA editing) (de Hoon et al. 2010; Li et al. 2018), 3’ adenylation or urydilation by terminal nucleotidyl transferases (Jones et al. 2009; Katoh et al. 2009), shortening of their 3’ end by poly(A)-specific ribonuclease (Lee et al. 2019), and, more rarely, shifts in their 5’ start sites (Tan et al. 2014). The diversity of miRNA isoforms (isomiRs) was initially proposed to increase the robustness of miRNA-mediated regulation, by fine-tuning the binding of miRNAs to their targets (Cloonan et al. 2011). Yet, there is now growing support for the notion that miRNA modifications may act as a conserved, additional layer of regulation of their activity (Fernandez-Valverde et al. 2010; Tan et al. 2014; Yu et al. 2017), as illustrated by the case of miR-222. Upon stimulation with interferon or *Salmonella*, shortening of the 3’ end of miR-222 occurs, and leads to a decreased apoptotic action of the miRNA, while maintaining an anti-proliferative effect through the binding of its canonical targets (Yu et al. 2017; Nejad et al. 2018). However, our understanding of the variability of isomiR expression across individuals and populations remains largely incomplete.

Following the canonical model, regulation of gene expression by miRNAs is achieved through the recognition of conserved target sites, which are mostly located in the 3’ UTR of protein-coding transcripts (Chi et al. 2009; Hafner et al. 2010; Kehl et al. 2017; Gebert and MacRae 2019). This binding typically results in the repression of the target protein by inducing mRNA deadenylation and degradation, or by inhibiting translation (Pasquinelli 2012; Gebert and MacRae 2019). Furthermore, a strong body of evidence highlights the importance of sequence complementarity between the miRNA seed region — located at position 2-7 from the 5’ end of the miRNA (Kehl et al. 2017; Gebert and MacRae 2019) — and its target site in determining miRNA-binding. Nonetheless, identifying which mRNAs are actively targeted by a given miRNA remains challenging (Khorshid et al. 2013; Gumienny and Zavolan 2015; Liu and Wang 2019). Previous studies of the regulatory impact of miRNAs on gene expression have reported conflicting results (Parts et al. 2012; Lappalainen et al. 2013; Siddle et al. 2014; Wang et al. 2018), possibly due to difficulties in disentangling the direct effects of miRNAs on mRNA degradation from co-transcription between miRNAs and their targets. In this context, RNA-seq can capture both steady-state gene expression levels, via the analysis of exonic reads, and the dynamic rate of transcription, through the quantification of intronic reads (Gaidatzis et al. 2015). In doing so, it offers a unique opportunity to determine the relative contribution of transcription and post-transcriptional regulation by miRNAs to gene expression variability.

In this study, we provide a comprehensive resource of genome-wide sequence-based miRNA diversity from primary human monocytes, both at the basal state and upon cellular treatment with four immune stimuli, originating from 200 individuals of African and European descent (100 individuals from each ancestry, Fig. 1). Leveraging the information obtained from 977 small RNA-sequencing profiles, together with whole-genome genotyping and exome sequencing data as well as mRNA-sequencing data from the same individuals, we define the levels of miRNA and isomiR diversity across individuals and populations, explore the genetic sources of miRNA expression variability and miRNA-environment interactions, evaluate the effects of immune challenges upon miRNA and isomiR expression dynamics, and quantify the relative impact of transcription and miRNA-mediated degradation on gene expression variability.

**Figure 1.**
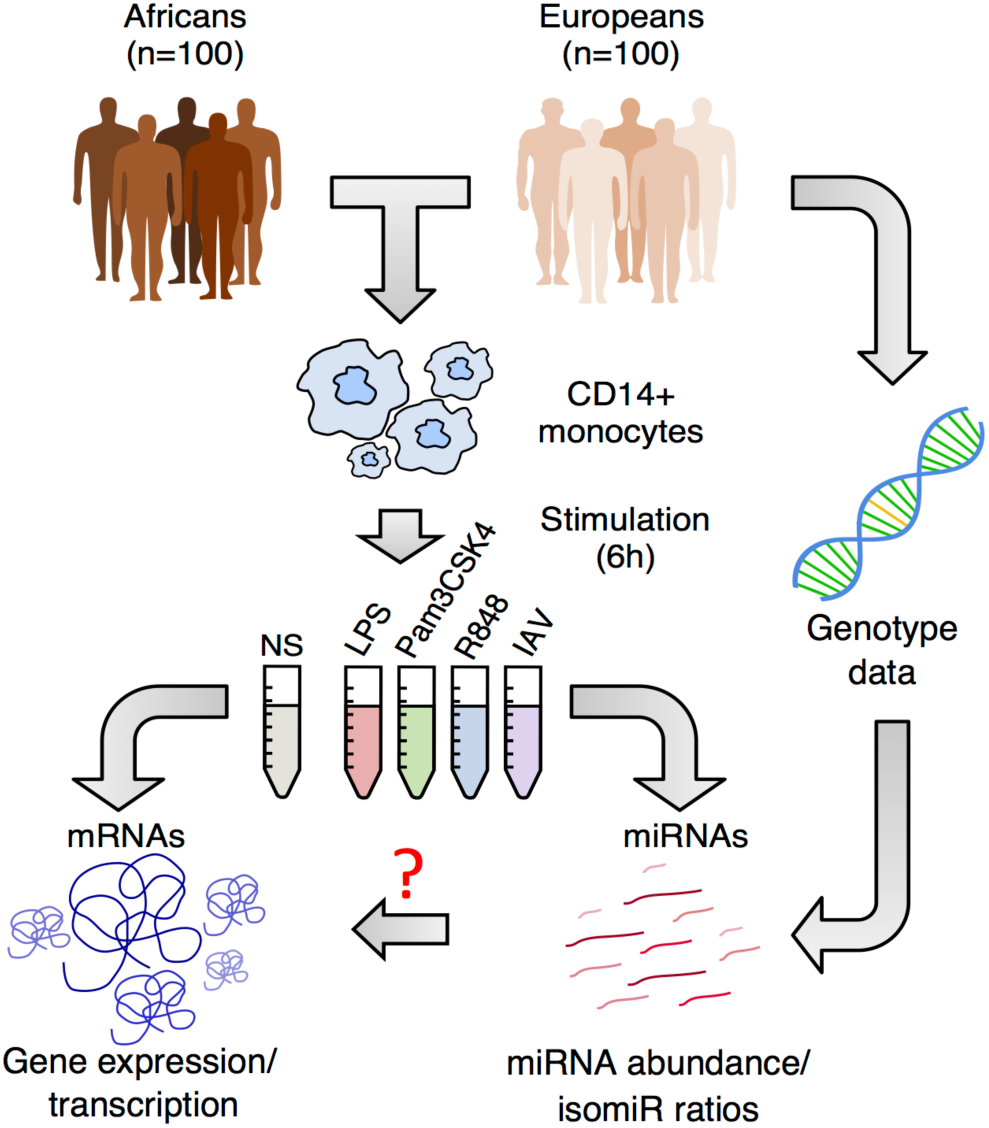
Population variation in miRNA response to immune activation. We stimulated monocytes from 200 healthy individuals using 3 TLR ligands as well as a strain of influenza A virus (IAV). For each individual, RNAs were extracted after 6h of stimulation. We sequenced small RNAs, from both stimulated cells and non-stimulated cells (NS) (this study). The integration of miRNA sequencing data with genetic data, obtained through whole genome genotyping, whole-exome sequencing and imputation (Quach et al. 2016), allowed us assessing the genetic bases of population differences in miRNA responses to stimulation, both quantitatively (miRNA abundance) and qualitatively (isomiR ratios). Furthermore, the availability of mRNA sequencing data from the same individuals and experimental conditions allows quantifying both total gene expression levels (exonic reads) and transcription rate (intronic reads derived from nascent mRNAs).

## RESULTS

### The landscape of miRNA and isomiR expression in human monocytes

We generated 977 small RNA sequencing profiles, in resting and activated cells, from 200 healthy individuals of African and European ancestry. Activation was performed for 6h with three different Toll-like receptor (TLR) ligands (LPS, Pam_3_CSK_4_ and R848 activating TLR4, TLR1/2 and TLR7/8 pathways, respectively), and a live strain of Influenza A Virus (IAV, Fig. 1). Small RNAs were separated from mRNAs, and sequenced at a mean depth of 12.4 million reads per sample (Methods; Supplemental Fig. S1A-C). After excluding reads outside the 18-26 nt range and low-quality samples (Supplemental Fig. S1D,E), we obtained an average of ~5 million reads aligned to miRNAs. To correct for cross-mapping artefacts between miRNAs, multiply-mapped reads were assigned to each possible locus using an Expectation-Maximization strategy (de Hoon et al. 2010). Library size was normalized across samples, and miRNAs with an average of <1 read per million miRNA-mapped reads (RPM) were discarded. This yielded a final set of 736 loci, encoding for 658 distinct miRNAs (Supplemental Table S1A).

Focusing on unique sequences, we identified 23,447 putative isomiRs, the vast majority (90%) of which were lowly abundant (<1 RPM; 14,277 isomiRs) or extremely rare (<1% of the reads of the associated miRNA; 6,811 isomiRs). Focusing on the remaining 2,359 unique miRNA sequences (corresponding to 492 loci encoding 451 distinct miRNAs, Supplemental Table S1B), we found that 86% of miRNAs expressed one or more isomiR(s) beside the canonical form, with a single miRNA expressing up to 8 frequent isomiRs (>5% reads) (Fig. 2A, Supplemental Fig. S2A,B). For more than 57% of miRNAs, the canonical isomiR accounted for less than half of the copies of the miRNA (Fig. 2B). Among the 311 miRNAs where the canonical isomiR was in minority (<50% of the reads), 25% had a seed sequence that differed from the canonical isomiR in more than 20% of their copies (Fig. 2C).

**Figure 2.**
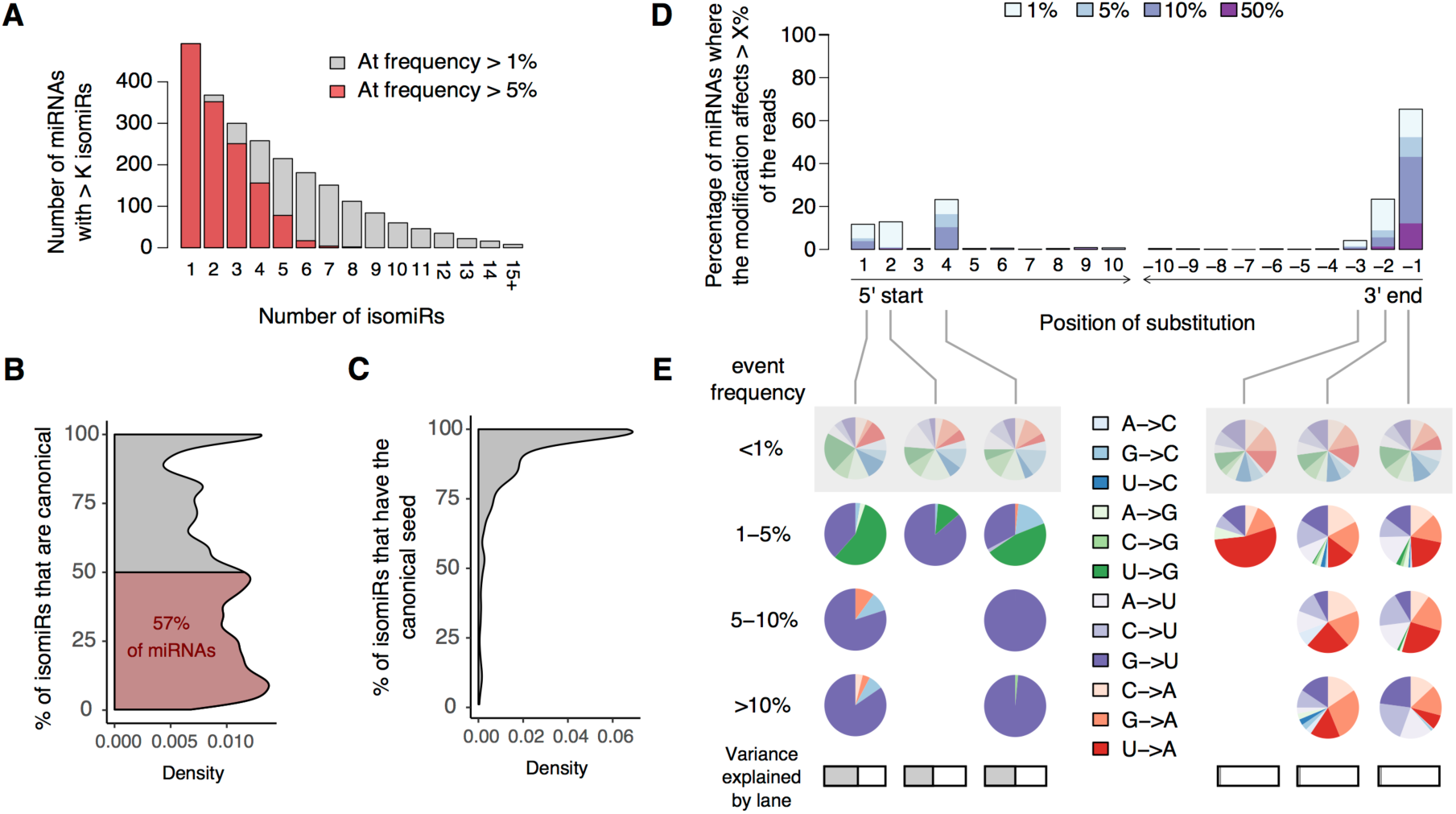
The landscape of miRNA diversity in human primary monocytes. (*A*) For each possible number of isomiRs *K*, number of miRNAs with more than *K* isomiRs at a frequency of >5% or >1%. (*B*) Distribution of the percentage of canonical isomiRs among all miRNAs; the area where the canonical isomiRs account for less than 50% of the reads is highlighted in red. (*C*) Distribution of the percentage of isomiRs with canonical seed among all miRNAs. (*D*) Distribution of edited nucleotides along miRNAs. For each nucleotide position, counting either from the 5’ start site (position 1 to 10) or from the 3’ end site (positions -10 to -1), we report the percentage of miRNAs that present an editing event accounting for ≥1% of reads (light blue). Similarly, we quantified the fraction of miRNAs where the editing event accounts for >5% (blue), >10% (indigo) or >50% (deep purple) of the reads. (*E*) Frequency of each type of substitution, according to the percentage of miRNA reads that are edited. Results are shown for positions where events were detected in >1% of miRNAs. At each position, distribution of substitution types among low frequency editing events (<1%), which we expect to be enriched in false positives, is provided as a reference (gray shadow). At each position, a horizontal grey bar indicates the variance in sample editing levels that is accounted for by the sequencing lane.

### Dissecting the mechanisms underlying miRNA isoform diversity

To dissect the processes leading to the high isomiR diversity observed, we pooled miRNAs across all conditions and quantified each type of miRNA modification separately (i.e. shifts in start/end site, non-template additions [NTA] and substitutions were quantified independently, and isomiRs with >1 modification were counted multiple times, see Methods). We found that shifts in 3’ end site of miRNAs were the most frequent type of modification, with 79% of miRNAs presenting a shift of their 3’ end site in > 5% of the reads, even after exclusion of non-template additions (Supplemental Fig. S2C,D, 87% including NTA), consistent with previous results (Siddle et al. 2015). Conversely, less than 32% of miRNAs presented a frequent shift of their 5’ start site (>5% of reads, Supplemental Fig. S2E), reflecting strong constraints on the miRNA seed.

Focusing on nucleotide substitutions, we found a strong enrichment of substitutions at 3’ end of the miRNA (binomial *p* < 3.7×10^−38^, Fig. 2D,E), recapitulating known patterns of 3’ terminal uridylation and adenylation (Ameres and Zamore 2013). We also detected a strong enrichment of substitutions at the 5’ end, as well as at the seed-altering positions 2 and 4 of the miRNA (binomial *p* < 2.2×10^−5^). All three positions (i.e. nucleotides 1, 2 and 4) presented a strong bias (binomial *p* < 9.8×10^−31^) toward G->U substitutions, as well as low frequency U->G changes at positions 1 and 4 (binomial *p* < 0.003). While the frequency of terminal substitutions was stable across sequencing batches (R^2^ < 0.1%), substitutions at positions 1, 2 and 4 depended on the sequencing lane (R^2^ > 48%), suggesting a technical bias. These substitutions were thus not considered for isomiR definition. IsomiR abundances were recomputed merging all isomiRs that differed by non-terminal substitutions and considering only isomiRs that result from shifts in the start or end of the miRNA or 3’ terminal uridylation and adenylation. After removing spurious isomiRs, the final dataset consisted of 2,049 common isomiRs across 435 miRNAs (Supplemental Table S1C).

We compared the frequency of miRNA modifications across both arms of the pre-miRNA hairpin (Supplemental Table S1D). We observed a stronger degree of 3’ terminal uridylation at -3p miRNAs (+12% of uridylated miRNAs on 3p arm compared to 5p; Wilcoxon *p* < 2.8×10^−11^, Supplemental Fig. S2F), consistent with the reported role of uridyl-transferases in pre-miRNA maturation (Heo et al. 2009; Kim et al. 2015; Kim et al. 2019). This increased uridylation was not associated to a higher rate of 3’ extensions among miRNAs located on the 3p arm (Wilcoxon *p* = 0.47), due to a higher rate of template extensions among 5p miRNAs (+7.6% on 5p arm compared to 3p; Wilcoxon *p* < 0.006, Supplemental Fig. S2G,H). Finally, we detected a higher usage of non-canonical, downstream start sites among -3p miRNAs (+3% compared to 5p miRNAs; Wilcoxon *p* < 0.003), consistent with a regulation of isomiR variability through the tuning of DICER positioning on the pre-miRNA (Zhu et al. 2018). Overall, the high variability of isomiRs detected highlights the complexity of the landscape of miRNA modifications in primary human monocytes.

### Marked effects of immune challenges upon miRNA and isomiR expression

Principal component (PC) analysis of miRNA abundances revealed a clear separation by stimulation conditions (Fig. 3A), after adjusting miRNA and isomiR expression for batch effects (date of experiment, date of library preparation and sequencing lane) and technical confounders (GC content and mean read length of the sample). PC1 opposed TLR-activated from IAV-infected samples, while PC2 captured the shared effect of all immune stimuli on gene expression. The variance explained by these PCs (i.e., a total of 16.7%) indicates that the effect of immune activation on miRNA expression is much weaker than that observed at the protein-coding mRNA level (Quach et al. 2016), where stimulation explained ~69% of expression variability. In contrast with patterns at the mRNA level, we noticed significant shifts between populations on both PCs (PC1, *t*-test *p* < 1.0×10^−79^; PC2, *t*-test *p* < 1.2×10^−11^), possibly reflecting differences in the intensity of miRNA responses to immune stimuli between individuals of African- and European-ancestry.

**Figure 3.**
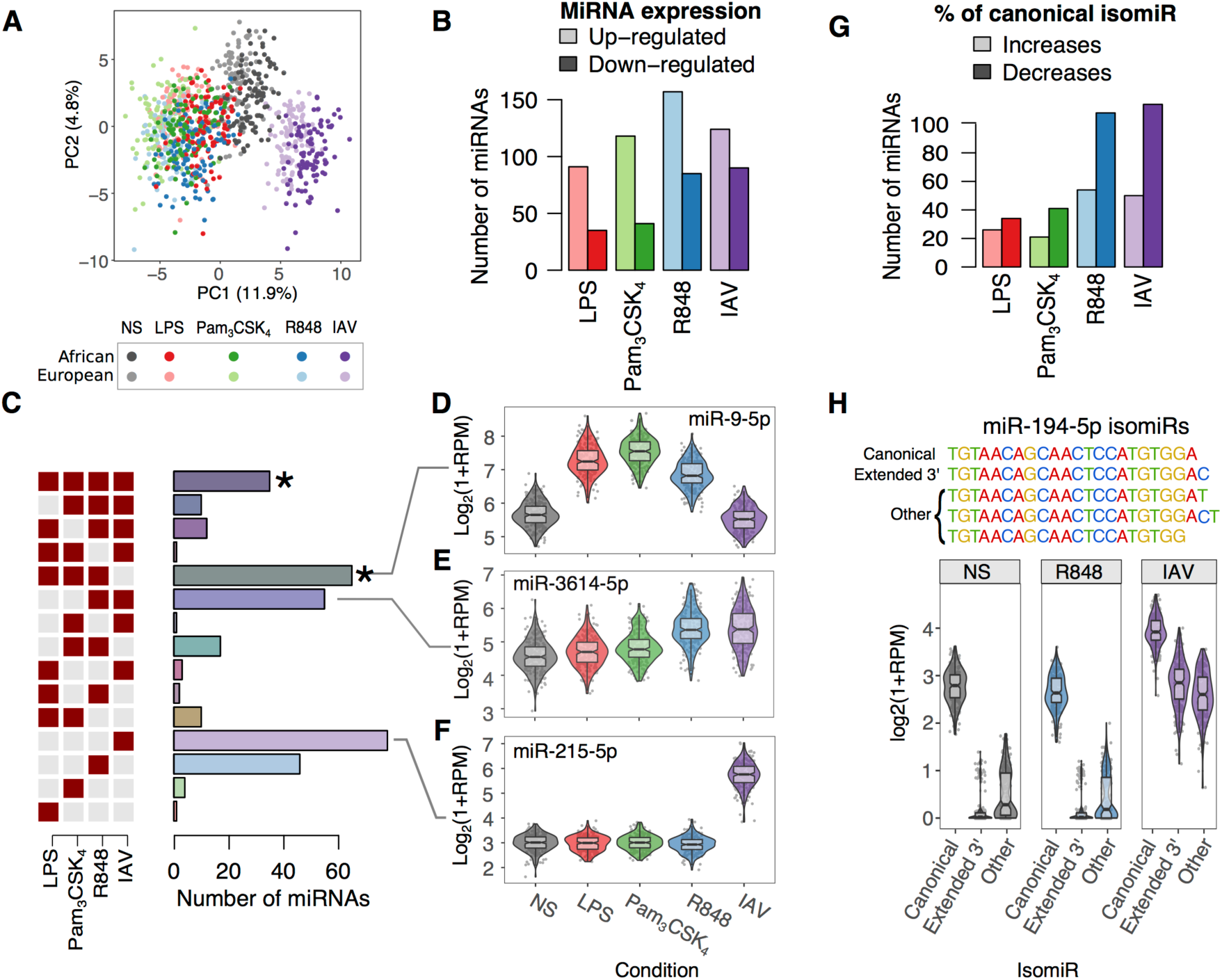
Stimulus-specific miRNA responses to immune activation. (*A*) PCA of log transformed miRNA abundances. Each dot represents a sample, colored according to the experimental condition (grey – non-stimulated, red – LPS, green – Pam_3_CSK_4_, blue – R848, purple – IAV). The same color code for conditions is used throughout the manuscript, and light and dark shades indicate European and African ancestry, respectively. (*B*) For each condition, number of DE miRNAs that are either up- (light shade) or down-regulated (dark shade). (*C*) Number of miRNAs that are differentially expressed (compared to NS) in a single condition or a combination of immune stimulations (*binomial *p* < 0.001, significance of overlap between stimuli). (*D-F*) Examples of DE miRNAs for the three most frequent patterns of differential expression across stimuli: (*D*) miR-9-5p, exhibiting a TLR-specific response. (*E*) miR-3614-5p, responding specifically to viral stimuli. (*F*) miR-215-5p, showing an IAV-specific response. (*G*) For each condition, number of miRNAs where the canonical isomiR is either up- (light shade) or down-regulated (dark shade). (*H*) Example of isomiR-levels response to IAV for miR-194-5p. miR-194-5p expresses 5 isomiRs that differ in their 3’ end. Violin plots show the expression of miR-194-5p isomiRs at the basal state, after R848 stimulation (as an example of a TLR-ligand), and IAV. The 3 least frequent isomiRs are grouped for clarity.

At FDR < 1%, we identified 340 miRNAs that presented differential expression upon stimulation (DE miRNAs, 30 with log_2_FC > 1), 233 of which were up-regulated in at least one condition (58-74% per condition; Fig. 3B; Supplemental Table S2A). Using a Likelihood-based Model Selection framework (Ding et al. 2018) (Fig. 3C), we estimated that 90% of DE miRNAs respond in a stimulus-dependent manner. The three most frequent patterns of miRNA responses were (i) a TLR-specific response (N = 65, 19% of DE miRNAs), as in the case of the NF-κB inhibitors miR-9-5p (Fig. 3D) and miR-155-5p, (ii) a viral-stimuli specific response (R848 and IAV, N = 55, 16% of DE miRNAs), such as miR-3614-5p recently involved in Crohn’s disease susceptibility (Wohlers et al. 2018) (Fig. 3E), and (iii) an IAV-specific response (N = 78, 23% of DE miRNAs), as attested by the pro-inflammatory mir-429 or the TRIM22 repressor mir-215-5p (Fig. 3F).

Focusing on how immune activation altered isomiR ratios, IAV infection clearly had the strongest impact (PC1, 4.7% of variance explained) followed by TLR7/8 activation (PC2, 2.6% of variance explained), with TLR4 and TLR1/2 activation showing a limited impact (Supplemental Fig. S3A). A total of 316 miRNAs changed their isomiR ratios upon stimulation (Supplemental Table S2B). Among these, the ratio of the canonical form was found to be affected for 212 miRNAs (67%), a ratio that decreases in 56% to 70% of the cases (Fig. 3G). For a majority of miRNAs, changes in isomiR ratios were of moderate intensity, with a single isomiR changing its ratio by more than 5% in only 5-11% of miRNAs (Supplemental Fig. 3SB). Notable exceptions included mir-155-5p, shifting toward extended 3’ isomiRs upon stimulation (13-33% increase in longer isomiRs, Wilcoxon *p* < 2.7×10^−68^), and miR-194-5p, showing an IAV-specific shift toward a 3’-extended isomiR (17% increase in longer isomiR upon infection, Wilcoxon *p* < 7.4×10^−97^, Fig. 3H). Overall, changes in isomiRs were most frequent following treatment with viral ligands (R848 and IAV), with 36% of isomiRs changes being shared between these stimuli (Supplemental Fig. S3C).

We next investigated whether this observation could be explained by specific mechanisms of miRNA modifications (Supplemental Table S2C). We found a significant trend toward a shortening of miRNAs at their 3’end site for both R848 and IAV conditions (Wilcoxon *p*_*adj*_ < 4.7×10^−14^, Supplemental Fig. S3D). This trend was observed for miRNAs located on both arms of their pre-miRNA hairpins, but was largely restricted to the IAV condition among 5p miRNAs (Wilcoxon test; *p*_*adj*_ = 0.017 and *p*_*adj*_ < 8.2×10^−24^ for R848 and IAV respectively, Supplemental Fig. S3D), suggesting a combination of multiple effects rather than a single mechanism. Specifically, 94% of the observed shifts in miRNA end sites were driven by changes of the miRNA reading frame rather than by a reduction of non-template additions. Yet, we observed significant changes in the frequency of 3’ end nucleotide additions, with increased adenylation and decreased uridylation (Wilcoxon test; *p*_*adj*_ < 2.2 ×10^−7^ and *p*_*adj*_ < 0.04, respectively), upon R848 and IAV stimulation (Supplemental Fig. S3E-F). Altogether, these results highlight significant shifts in miRNAs expression upon immune stimulation, and reveal a high rate of isomiR modifications in response to viral stimuli.

### Strong selective constraints limit miRNA expression variability

To assess the extent to which miRNA expression variability is under genetic control, we focused on the 598 miRNAs associated to a unique genomic location and searched for genetic variants associated with changes in miRNA abundances within a 1Mb window around each miRNA (miRNA Quantitative Trait Loci, or miR-QTLs). At 5% FDR, we identified 122 miRNAs associated with at least one miR-QTL (Supplemental Table S3A), corresponding to ~20% of the tested miRNAs. Interestingly, this proportion was lower than that observed for mRNAs of protein-coding genes or long non-coding RNAs (with 31% and 33% of the latter presenting an eQTL, respectively, Fisher’s *p* < 1.2×10^−7^). However, we found a comparable proportion of eQTLs among transcription factors and loss-of-function intolerant genes (25% and 22% of eQTLs, respectively, Fig. 4A). Furthermore, we observed a decreased proportion of miR-QTLs among highly expressed miRNAs (Supplemental Fig. S4A,B). Interestingly, miRNAs with conserved promoters (mean phastCons > 20%) were depleted in miR-QTLs with respect to miRNAs with less conserved promoters (OR = 0.54, Fisher’s *p* < 0.008). In addition, miR-QTLs of miRNAs with a conserved promoter were located on average further away from the transcription start site (TSS; + 3.5 kb, Wilcoxon *p* < 0.03, Supplemental Fig. S4C). Collectively, these observations indicate strong selective constraints that limit miRNA expression variability.

**Figure 4.**
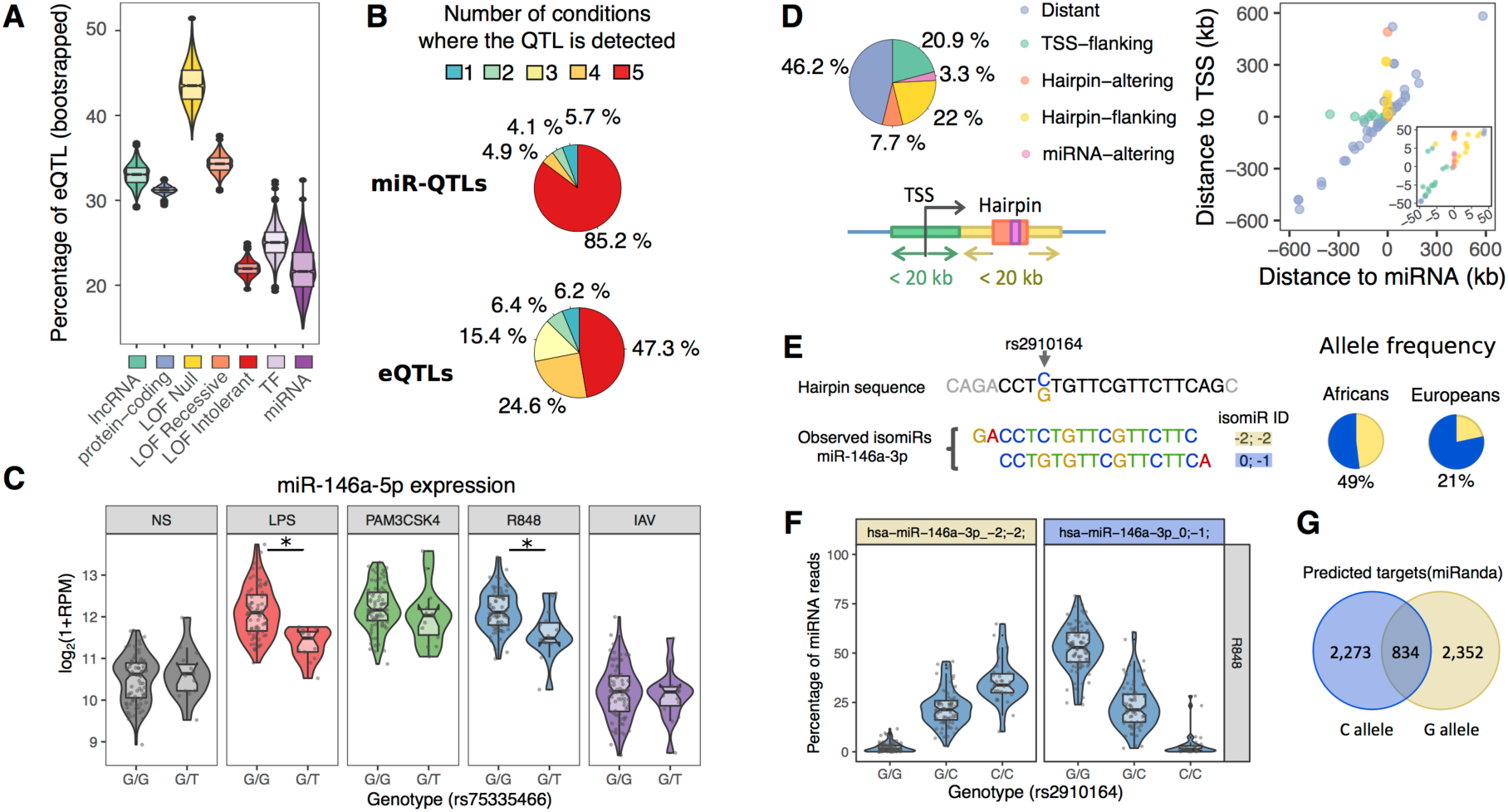
Genetic basis of miRNA expression upon immune activation. (*A*) Percentage of genes and miRNAs under genetic control, across various functional classes. For each gene class, 1000 bootstrap resamples were performed and the resulting distribution is shown as a boxplot. (*B*) Sharing of miR-QTLs and eQTLs across conditions. For the 122 miR-QTLs and 3,802 eQTLs, number of conditions where a QTL is detected. (*C*) Example of an African-specific response miR-QTL. The rs75335466-T allele is associated with reduced expression of miR-146a-5p specifically upon stimulation by LPS and R848 (* *t*-test *p-value* < 0.001). For clarity, data is shown only for African-ancestry individuals. (*D*) Localization of miR-QTLs. Left: Frequency of miR-QTLs that either overlap the mature miRNA (*miRNA-altering*, pink) or its hairpin (*hairpin-altering*, orange), or are located <20kb away from the miRNA hairpin (*hairpin-flanking*, yellow) or TSS (*TSS-flanking*, green). Remaining miR-QTLs are annotated as *Distant* (blue). Right: Distance of mir-QTLs from the mature miRNAs (*x*-axis) and its associated TSS (*y*-axis). Each miR-QTL is shown as a separate dot, colored according to its localization. Negative distance indicates that the miR-QTL is located upstream of the miRNA and/or TSS. Close up view is shown for miR-QTLs located < 50 kb from the miRNA or TSS. (*E-G*) Impact of rs2910164 variant on miR-146a-3p isomiRs. (*E*) Genomic context and frequency of the rs2910164 variant. The rs2910164 C/G variant is shown with its neighboring hairpin sequence. Sequence of the canonical miRNA is displayed in black. The 2 most common isomiRs of miR-146a-3p are displayed below and denoted as (−2; −2), and (0; −1) based on the coordinates of their start/end site relative to the canonical miRNA sequence. Note that the C/G substitution is not considered for the quantification of (−2; −2) and (0; −1) isomiRs. Frequency of C/G alleles in our sample is shown in African- and European-ancestry individuals separately. (*F*) isomiR-QTL of miR-146a-3p. Ratios of (−2;−2) and (0; −1) isomiRs are shown for each genotype, in the R848 condition where the isomiR-QTL is the strongest. For clarity, other isomiRs are not displayed. (*G*) Overlap of miRNA targets predicted by miRNA for each possible isomiRs.

### miR-QTLs are largely shared across immune stimuli

When comparing the occurrence of miR-QTLs across stimuli, we found that 85% of all miR-QTLs were shared across all experimental conditions (Fig. 4B; Supplemental Table S3B), with a minority (N=18, 15%) displaying condition-dependent effects and only 5.7% being specific of one condition. This observation is in stark contrast with eQTLs of protein-coding genes, where 53% of them displayed condition-dependent effects (Fig. 4B). However, the proportion of miR-QTLs and eQTLs that are specific of one condition was quite similar (5.7% and 6.2% respectively; Fig. 4B). Among condition-dependent miR-QTLs, we detected 4 response miR-QTLs, i.e., genetic variants that manifest their effects on miRNA abundance only in the presence of immune stimulation (*p*_interaction_< 0.001, corresponding to 5% FDR and not detected in the non-stimulated state). For example, the African-specific rs75335466 has a derived allele (Derived Allele Frequency (DAF) = 7.5% in African-ancestry individuals) that is associated, upon stimulation of TLR4 and TLR1/2 pathways, to a reduced up-regulation of the dominant arm of miR-146a (miR-146a-5p, *p*_interaction_ < 5.6×10^−4^, Fig. 4C), which acts as an inhibitor of TRAF6 and IRAK1 (Taganov et al. 2006).

Given the reported role of enhancers in leading responses to immune stimulation (Arner et al. 2015), we hypothesized that the limited condition-specificity of miR-QTLs might be driven by a different regulatory architecture from that of eQTLs. We first sought to characterize the regulatory elements underlying miRNA expression. We found that 54% of miR-QTLs were located < 20kb from either the TSS of the pri-miRNA they regulate or the pre-miRNA hairpin that contains the mature miRNA (Fig. 4D). Furthermore, we observed a strong over-representation of both promoters (OR=17, Fisher’s *p* < 4.5×10^−12^) and enhancers (OR=3.6, Fisher’s *p* < 5.9×10^−5^) among miR-QTLs. Yet, the percentage of miR-QTLs falling into promoters or enhancers did not differ significantly from that observed for eQTLs (Fisher’s exact test; *p* = 0.25 for promoters and *p* = 0.88 for enhancers; Supplemental Fig. S4D). Overall, these results reveal that despite a similar promoter/enhancer architecture, miR-QTLs are largely insensitive to stimulation compared to eQTLs of protein-coding genes.

### Limited genetic control of isomiR diversity

We then searched for genetic variants that alter isomiR ratios (isomiR-QTLs). Only 25 isomiRs were associated with at least one isomiR-QTL, involving 13 miRNAs (Supplemental Table S3C), 84% of these being shared across conditions. Note that because we did not consider non-terminal substitutions in our definition of isomiRs, these numbers do not take into consideration genetic variants that directly alter the miRNA sequence, unless they also alter the start/end site of the miRNA. An interesting case of isomiR-QTL is provided by the rs2910164 variant (DAF: 49% in African and 21% in European ancestry groups), which disrupts the seed of the passenger arm of miR-146a (miR-146a-3p, Fig. 4E). The derived allele of rs2910164 (G) is associated with both an increase in expression of miR-146a-3p (*t*-test, |β_miR-QTL_| > 0.31, *p* < 3.1×10^−7^) and a shift of both the start and end sites of the mature miRNA (*t*-test, |β_isomiR-QTL_| > 0.15, *p* < 2.1×10^−9^, Fig. 4F). This shift leads to a complete redefining of the miR-146a-3p targets, with 2,273 predicted targets being lost (73%) and 2,352 novel targets being gained (Fig. 4G). Interestingly, the rs2910164 G allele is associated with increased risk of allergic rhinitis (*p* < 1.9×10^−13^) and asthma (*p* < 6.2×10^−9^) in the GWAS Atlas (Watanabe et al. 2019). Despite the limited genetic control of isomiR diversity, these results suggest that genetic variants altering isomiR ratios have a significant impact on immune response variability.

### Marked differences in miRNA expression related to population ancestry

We subsequently explored the extent to which miRNA responses to stimulation differ between individuals of African and European ancestry. We identified a total of 351 miRNAs whose transcriptional profiles differed between populations in at least one experimental condition, either in abundance (pop-DE-miR, N=244, including 141 with |log_2_FC|>0.2, Fig. 5A and Supplemental Table S4A), or in isomiR ratios (pop-DE-isomiR, N=188, including 148 with Δ_isomiR-ratio_ > 1%, Fig. 5B; Supplemental Table S4B), with 81 miRNAs differing in both expression and isomiRs. We found that at the basal state population differences in expression of miRNAs were similar in magnitude to those of protein-coding genes (17% of miRNAs and 12% of protein-coding genes display a |log_2_FC|>0.2 between populations, Wilcoxon *p* = 0.08, Fig. 5C). Upon stimulation, however, protein-coding genes displayed a marked increase in population differences (17%-26% with a |log_2_FC|>0.2 between populations, Wilcoxon *p* < 2.7×10^−27^), while population differences in miRNA expression remained rather stable (14-18% with a |log_2_FC|>0.2 between populations, Wilcoxon *p* > 0.26, Fig. 5C).

**Figure 5.**
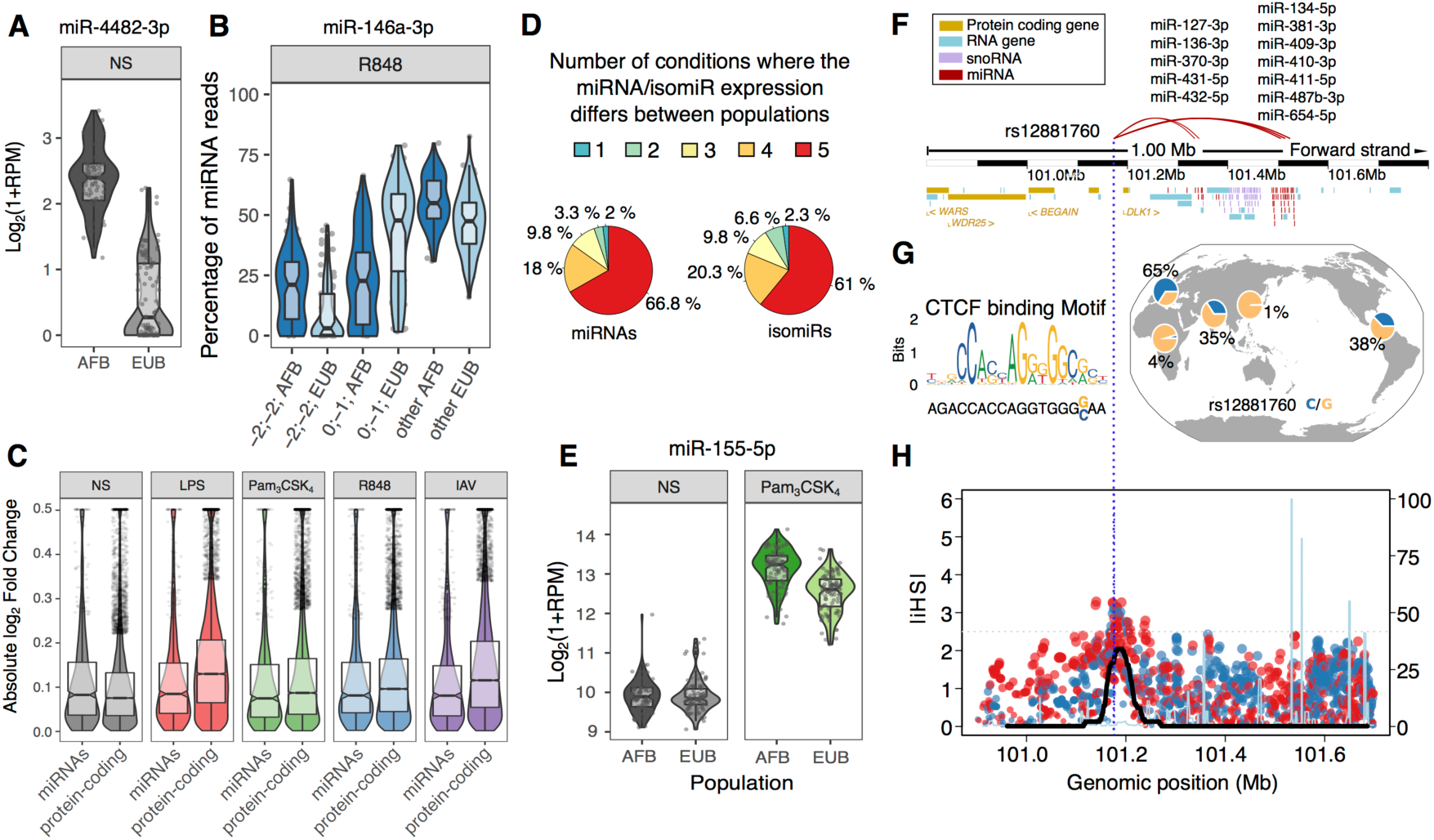
Population differences in miRNA expression. (*A*) Example of a miRNA (miR-4482-3p) differentially expressed between populations. Expression of miR-4482-3p is shown separately for African- (AFB) and European- (EUB) ancestry individuals. (*B*) IsomiRs of miR-146a-3p are differentially expressed between populations. For each population and isomiR, isomiR ratios are shown in the R848 condition where the difference is the strongest. All isomiRs with <1 RPM on average, are pooled and annotated as *other*. (*C*) Amplitude of population differences in expression of miRNAs and protein-coding genes, at basal state and upon stimulation. (*D*) Sharing of pop-DE-miRs and pop-DE-isomiRs across conditions. For the 244 pop-DE-miRs and 188 pop-DE-isomiRs, number of conditions where we observed a difference between populations. (*E*) Expression of miR-155-5p is differential between African- and European-ancestry individuals, specifically in TLR-stimulated conditions. For simplicity, only Pam_3_CSK_4_ condition is shown. (*F-H*) Signatures of positive selection targeting the miR-QTL hotspot rs12881760. (*F*) Genomic context of the miR-QTL hotspot, displaying protein coding genes (yellow), RNA genes (cyan), snoRNAs (purple) and miRNAs (red) in a 1Mb-window around the locus. Red lines link the miR-QTL to its target miRNAs, the name of which are indicated above. (*G*) Impact of the rs12881760 variant on a CTCF motif, and worldwide frequency of the motif-disrupting C allele. (*H*) Signatures of positive selection at the rs12881760 locus. |iHS| are displayed for all SNPS with MAF > 5% in Europeans, and dots are colored according to the sign of the iHS statistic (red: positive; blue: negative). The black line indicates the percentage of outliers (|iHS|>2.5) on a sliding window of 100 consecutive SNPs with a MAF>5% (right axis). Recombination rate is overlaid in light blue and normalized to the maximum recombination rate in the region (peak:152 cM/Mb).

Population differences in miRNA expression and isomiRs were largely shared across experimental conditions, with 67% of popDE-miR and 61% of pop-DE-isomiRs being shared across all experimental conditions (Fig. 5D). Yet, we identified 9 miRNAs that displayed population differences only upon stimulation (Supplemental Table S4C), including key immune modulators such as the pro-inflammatory miR-155-5p, which showed marked population differences upon TLR1/2 stimulation (Pam_3_CSK_4_; *p*_interaction_ < 1.0 ×10^−9^ Fig. 5E). Looking at the rate of miRNA modifications, we found that the 3’-end shortening of miRNAs located on the 3p-arm was more frequent in individuals of African ancestry with respect to those of European-ancestry (Wilcoxon p < 3.2×10^−6^ at basal state, Supplemental Table S4D). Furthermore, individuals of African ancestry presented, upon stimulation, an increased rate of 3’ adenylation, regardless of the arm where the miRNA is located (Wilcoxon *p* < 4.5×10^−5^), partially compensating the detected shortening of miRNAs located on the 3p arm.

### Sources of ancestry differences in miRNA and isomiR responses

We next searched for the sources of population differences in miRNA expression, and found a significant enrichment of popDE-miR and -isomiR in miRNAs whose expression is under genetic control (i.e., miR- or isomiR-QTL; OR > 1.7, Fisher’s *p* < 1.1×10^−2^). By computing the fraction of population differences in miRNA expression that is attributable to genetic factors, we estimated that, among the 57 popDE-miR with a miR-QTL (23% of popDE-miR), genetics accounted for ~60% of population differences on average. Across all miR-QTLs, the strongest differences in frequency between individuals of African and European ancestry was observed at the variant rs12881760 on chromosome 14. This variant is associated with the expression of 12 miRNAs that are located in a cluster of 148 small RNAs spanning over 250kb (Fig. 5F). The derived allele (C) disrupts a CTCF binding site located ~200kb upstream of the small RNA cluster, and is associated with a lower platelet mass in the GWAS Atlas (p < 6.5 × 10^−48^) (Watanabe et al. 2019). Interestingly, the C allele is found at high frequency in European-descent populations (e.g., up to 72% in Iberians) and rare in Africans and East Asians (<4%, Fig. 5G). Moreover, it harbors a strong signature of positive selection in Europe (iHS = −3.10, *p*_emp_=0.002, 31% of SNPs with |iHS|>99^th^ percentile in a 100 SNP window around the locus, *p*_enrich_=0.003, Fig. 5H), clearly supporting a history of recent adaptation targeting this locus. Overall, while a substantial fraction of population differences may be due to non-genetic factors, our results show that genetic differentiation at miR-QTLs has, in some cases, substantially contributed to population differences in miRNA expression.

### The regulatory impact of miRNA variability on downstream immune responses

Finally, we quantified the extent to which miRNAs contribute to the regulation of immune-related gene expression. To do so, we leveraged mRNA sequencing data obtained for the same individuals, and correlated miRNA expression with mRNA levels of 12,578 genes expressed in our monocyte setting (FPKM>1) (Quach et al. 2016), using stability selection (Methods). At an 80% probability threshold (~1% FDR based on permutations, Supplemental Fig. S5), we found that 25-45% of genes were significantly associated with at least one miRNA, with a single gene being independently associated with up to 6 different miRNAs per condition (Fig. 6A). Among conditions, the number of genes associated with a miRNA was slightly higher for viral stimuli (39-44% for R848 and IAV vs. 24-33% for NS, LPS and Pam_3_CSK_4_, Fig. 6A). Surprisingly, among the 6,009 miRNA-gene associations detected at the basal state, only 43% displayed negative associations, of which 12% presented a known binding site for their associated miRNA. In addition, we found that predicted miRNA targets were depleted in negative correlations with their cognate miRNAs (OR = 0.83, Fisher’s *p* < 0.02). These results suggest that, at least in our setting, miRNA-driven transcript degradation has a minor impact on gene expression variability, with correlations being likely explained by co-transcription of miRNAs and their target genes.

**Figure 6.**
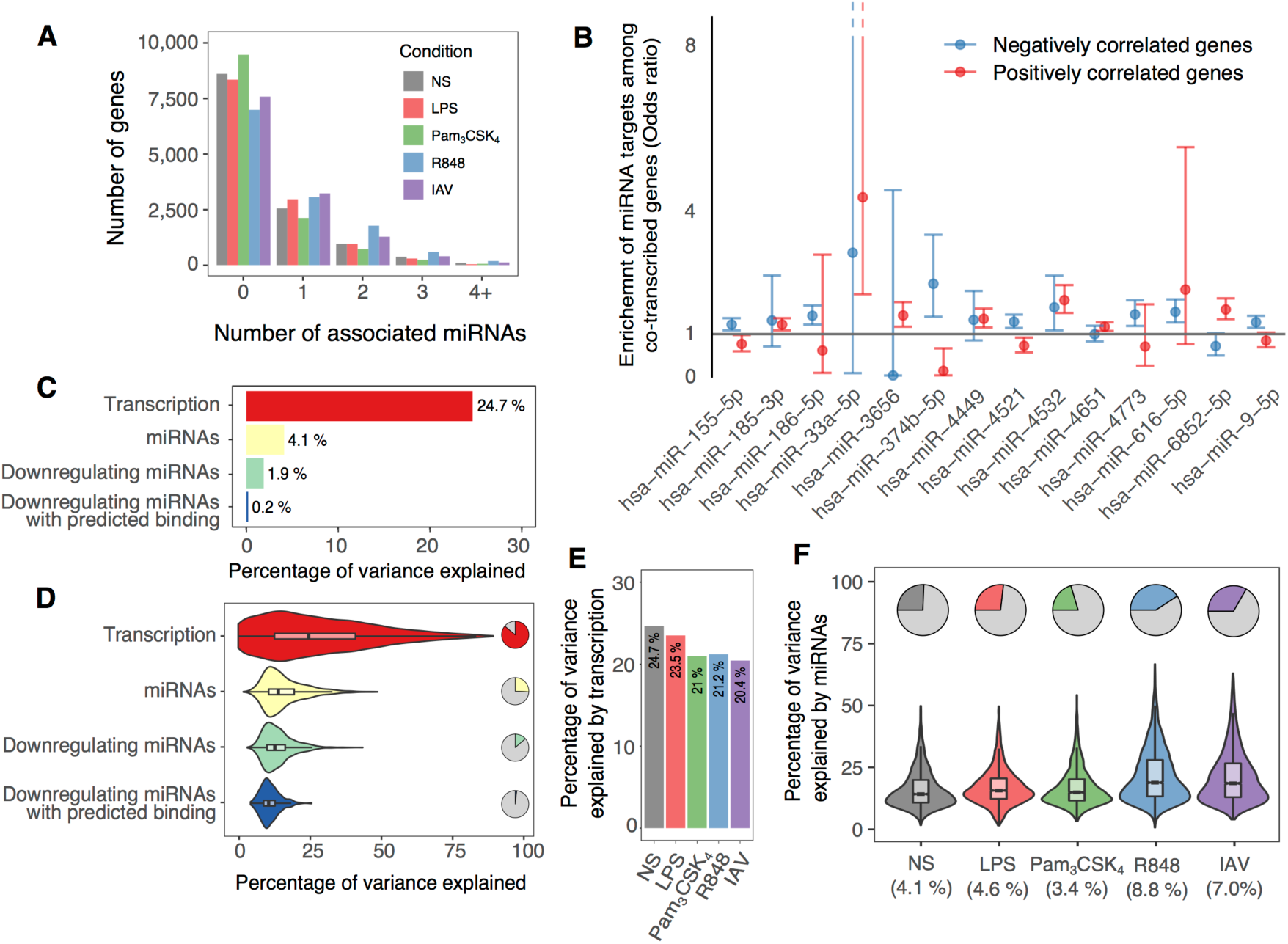
Impact of miRNA levels on gene expression. (*A*) Distribution of the number of associated miRNAs per gene according to the experimental condition. (*B*) Enrichment or depletion of miRNA targets among genes whose transcription level correlates with miRNA expression (co-transcribed genes). For each miRNA, odds ratios are reported separately for genes whose transcription is positively (red) or negatively (blue) correlated to miRNA expression. Enrichments are displayed only for miRNAs that have a significant enrichment of their targets in either positively or negatively correlated genes (5% FDR). (*C-D*) Percentage of gene expression variance that is attributable, at the basal state, to either transcription or miRNA variation. For miRNAs, attributable variance is also reported considering only negative associations, or negative associations with predicted binding between the gene and the miRNA. (*C*) Global percentages (average values across all genes). (*D*) Distribution at the gene level. Pie charts indicate the percentage of genes associated with transcription or miRNAs. Violin plots display the distribution of the variance attributable to each factor, among significantly-associated genes. (*E*) Global percentage of variance attributable to transcription according to the experimental condition. (*F*) Within each experimental condition. pie charts indicate the percentage of genes associated to at least one miRNA. Violin plots display the variance attributable to miRNAs among significantly-associated genes.

To test this hypothesis, we quantified intronic reads that derive from unspliced, nascent transcripts, as a measure of transcription rate (Gaidatzis et al. 2015). We identified widespread miRNA-gene correlations, each miRNA correlating with transcription rates of ~100 genes at the basal state (min:1, max: 2,680) and up to 173 upon stimulation (min:1, max: 4,137). We found 7 miRNAs that are co-transcribed with their target genes, i.e., enrichment of miRNA targets in *trans* among genes positively correlated at the transcription level (Fig. 6B). Among these, the regulator of cholesterol homeostasis miR-33a-5p (OR=4.3, Fisher’s *p* < 1.7×10^−4^) balances the effect of its host gene, the TF *SREBF2*, on fatty acid synthesis/uptake by repressing the cholesterol transporter ABCA1 (Najafi-Shoushtari et al. 2010). We also identified 7 miRNAs negatively correlated to the transcription of their target genes (Fig. 6B), suggesting a feed forward loop mechanism, where miRNA down-regulation occurs in parallel to the transcription of its target genes to promote rapid expression changes. These include key regulators of the immune response such as the NF-κB inhibitors miR-9-5p (OR=1.2, Fisher’s *p* <5.8×10^−4^) and miR-155-5p (OR=1.3, Fisher’s *p* <1.0×10^−5^).

Using a variance partitioning approach, we quantified the amount of inter-individual variation in gene expression attributed to either gene transcription or miRNA expression (Zuber and Strimmer 2011). We found that, on average, transcription accounts for 25% of the variance in gene expression at the basal state, with this amount decreasing upon stimulation (min: IAV-20%, max LPS-24%, Wilcoxon *p* < 2.8×10^−3^, Fig. 6C-E). Conversely, miRNA expression variability accounted for only 4.1% of the total variance of expression of their associated genes, and between 3.4% (Pam_3_CSK_4_) and 8.8% (R848) upon stimulation (Fig. 6C,D,F). These figures decreased to ~0.2% when focusing only on negative associations, and disregarding miRNAs with no predicted targets for the gene under consideration (Fig. 6C,D). Testing for the downstream effects on gene expression of the 122 miR-QTLs and 25 isomiR-QTLs, we found no evidence for an enrichment of *trans*-effects compared to random SNPs matched for allele frequency (π_1_^obs^ = 0.5%, resampling *p*=0.36). While miRNAs display significant correlations with gene expression, our results overall indicate that miRNA variation has a limited impact on gene expression variability.

## DISCUSSION

Several important insights can be drawn from our study. First, we show that upon immune stimulation or infection the miRNA repertoire is subject to important modifications that are not only quantitative, through the modulation of miRNA expression, but also qualitative, through changes in isomiR proportions (Siddle et al. 2015; Nejad et al. 2018). Although isomiR modifications can be confounded by cross-mapping artifacts and sequencing errors (de Hoon et al. 2010), we reduced here the impact of such technical biases by focusing on frequent, biologically-plausible modifications and excluding those that correlate with technical covariates. In doing so, we detected systematic shifts in isomiR proportions occurring primarily upon cellular treatment with viral challenges. While most changes in isomiR usage observed at 6h of stimulation are of modest effect size, it is possible that they anticipate more drastic modifications occurring later in time, as shown in the case of bacterial infections (Siddle et al. 2015). Whether induced by stimulation or associated with genetic variants, changes in isomiR usage have the potential to deeply alter miRNA-gene interactions (Yu et al. 2017), which may, in turn, impact organismal phenotypes. This is illustrated by miR-146a-3p, where a genetic variant inducing a shift in miRNA boundaries that leads to a broad redefinition of its targets, has been associated with immune pathologies. Our work highlights the importance of considering isomiR changes, and not only miRNA expression, when studying the impact of miRNAs on immune responses (Yu et al. 2017; Nejad et al. 2018).

Several lines of evidence indicate that strong selective constraints limit miRNA variability in response to immune challenges. First, we observe that both the response of miRNAs to immune activation and its variability between populations is much more nuanced than that of protein-coding genes. Second, we report a limited number of miRNAs, as well as isomiRs, whose expression levels are associated with miR-QTLs and isomiR-QTLs, even among highly expressed miRNAs. Finally, in contrast with protein-coding eQTLs, which are largely context-dependent and thus variable across stimuli, we find that the detected miR-QTLs are mostly unaffected by immune stimulation. Our analyses thus support the notion that the miRNA system is little tolerant to genetic variation modulating its response to stimulation.

Despite the global constraints driving miRNA variability, we find marked differences in miRNA expression between individuals of African and European ancestry, with ~25% of miRNAs per condition being differentially expressed between populations. The limited genetic control of miRNA expression, with respect to protein-coding genes, indicates that non-genetic factors may play an important role in the observed population differences. While over 60% of expression differences related to ancestry are unaffected by stimulation, we identified 9 miRNAs that present population differences uniquely upon immune challenge. Of these, 5 have been previously reported to correlate with LPS tolerization in mice (Seeley et al. 2018), including key immune regulators such as miR-155-5p and miR-222-3p. This suggests that population differences in miRNA responses are driven by specific environmental exposures, possibly through epigenetic priming of innate immune response.

Despite non-genetic factors plays a preponderant role in explaining ancestry-related differences in miRNA expression, genetic factors can also significantly contribute to population differences; genetics explains ~60% of the observed differences among miRNAs associated with a miR-QTL. In some cases, population differences driven by genetic factors can be differently distributed between African- and European-descent groups. For example, the frequency of the variant rs290164 is sufficiently different between populations (ΔDAF = 28%) to explain ~76% of the differences in isomiR ratios for miR-146a-3p. Furthermore, we identify a European-specific variant (rs12881760), which controls a cluster of 12 miRNAs in *cis*, that displays extreme population differentiation (*F*_ST_) with Africans (top 0.2% of *F*_ST_) and East Asians (top 0.004% of *F*_ST_). The adaptive nature of this variant in Europeans is supported by a strong enrichment of |iHS| outliers at this locus that, moreover, has been associated with platelet parameters, likely underlying differences in platelet activity between European- and African-ancestry individuals (Edelstein et al. 2013). Interestingly, an independent event of positive selection targeting the same miRNA-rich cluster has been detected in Asian populations (Quach et al. 2009), highlighting the adaptive role of this locus in populations of non-African ancestry.

Finally, our study-design allowed us to assess the relative contribution of transcriptional regulation and miRNA-mediated degradation on downstream immune responses. While we find a strong effect of transcription rate on gene expression, our model predicts that miRNA-mediated degradation accounts for <0.2% of the variation in gene expression, suggesting a limited effect of miRNAs on mRNA stability. Such a modest impact is further supported by the lack of measurable effects of miR-QTLs on gene expression, at least in miRNAs that tolerate genetic variation (miR-eQTLs). Although these results are at odds with the canonical model of miRNA activity (Pasquinelli 2012), they are consistent with the reported low levels of miRNA-mRNA correlations and the frequent occurrence of positive correlations between miRNA expression and that of their predicted targets (Rantalainen et al. 2011; Parts et al. 2012; Lappalainen et al. 2013; Siddle et al. 2014). Our study provides a model that could explain the occurrence of such positive correlations, as attested by cases where miRNA expression is correlated with the transcription of their targets, creating either feedback loops, as for miR-33a-5p, or feed-forward loops, as for the TLR-induced miR-9-5p and miR-155-5p. When adjusting for transcription rate, miRNA expression captures 3 to 6% of the variation in gene expression on average. This could reflect either an indirect contribution of miRNAs to immune response variability through translation inhibition of key immune regulators, or residual co-transcription due to the dynamic nature of gene transcription.

Together, this study shows that genetic and non-genetic factors contribute to marked population differences in miRNA abundance and isomiR ratios. Yet, we also show that such differences have a moderate impact on the transcriptional landscape of immune cells at 6h, suggesting that the consequences of miRNA deregulation may be most visible at later stages of the immune response. Overall, our study reports a large set of miRNAs and isomiRs that present differential responses across bacterial- and viral-like challenges and/or between populations of different ancestry during an early time window of the innate immune response. In doing so, it constitutes a useful resource for evaluating the role of these epigenetic regulators in host differences of immunity to infection and susceptibility to disease, both at the individual and population levels.

## METHODS

### Ethics statement

Human primary monocytes were obtained from healthy volunteers who gave informed consent. All experiments were approved by the Ethics Board of Institut Pasteur (EVOIMMUNOPOP-281297) and the relevant French authorities (CPP, CCITRS and CNIL).

### Samples and dataset

Biological samples were generated as part of the EvoImmunoPop project (Quach et al. 2016). Briefly, the EvoImmunoPop cohort is composed of 200 healthy, male participants of self-reported African and European descent, recruited in Belgium (100 individuals of each population). For all individuals, total RNAs from CD14-positive cells treated for 6 hours with five conditions of stimulation (resting, LPS, Pam_3_CSK_4_, R848, and IAV A/USSR/90/1977). Genotyping was performed using both Illumina HumanOmni5-Quad BeadChips and whole-exome sequencing with the Nextera Rapid Capture Expanded Exome kit. Stringent quality control and imputation procedures were applied (Quach et al. 2016), leading to a final set of 19,619,457 SNPs, of which 9,854,620 SNPs had a minor allele frequency (MAF) greater than 5% in either population of our cohort. Regarding the mRNA sequencing dataset, libraries from total RNA samples and transcriptome sequencing were performed using TruSeq RNA Sample Prep Kit v2 for mRNA library construction, TruSeq SR Cluster Kit v3-HS for cluster generation, and TruSeq SBS kit v3-HS for sequencing on an Illumina HiSeq2000 platform. In total, an average of 34.4 million 101-bp single-end reads per sample (min: 27.7-max: 94.8 million reads) were obtained (Quach et al. 2016). High-density genotyping and exome sequencing data, and the mRNA sequencing data, used in this study are available in the European Genome-Phenome Archive (EGAS00001001895).

### Small-RNA Library preparation and sequencing

Total RNA samples have been used to generate miRNA sequencing data. Low molecular weight RNA fragments were selected by gel excision (targeting fragments of ~22 bp), and sequencing libraries were prepared using the Illumina TruSeq small RNA library prep Kit. Indexed cDNA libraries were then pooled by groups of 18 (in equimolar amounts), and sequenced with single-end 50bp reads on the Illumina HiSeq2000. After exclusion of one sample that yielded less than 1.8 million read counts, we obtained an average of 12.4 million raw reads per sample with a minimum yield of 8.0 million reads.

### Pre-processing of raw sequencing reads

Sequences matching the 3’ adaptor sequence were identified and trimmed, using fastx_clipper version 0.0.13 with the following options –l 0 –n –M 10, to require a minimum adapter alignment length of 10 base pairs, while keeping all sequences regardless of their length or presence of unknown nucleotides. This led to the exclusion of ~2% of reads per sample, and final read lengths ranged from 1 to 42 bases. We confirmed that all samples had average base quality (Q) values >30 at all positions, and that per-base GC distributions were within expected ranges. We further checked that read length distributions showed an enrichment of ~22 bases-long reads for all samples, consistent with expectations for mammalian miRNAs (~22 bases), discarding reads shorter than 18 or longer than 26 bases. After these filtering steps, we obtained an average of 8.8 million (minimum 4.1 million) short reads per sample that were used for small RNA quantification.

### Sequence alignment

Sequences were aligned to the human reference genome (build GRCh37/hg19) using bowtie (version 1.1.1) (Langmead et al. 2009). We mapped reads allowing for 2 mismatches (-v 2) and reported all best alignments for reads that mapped equally well to more than one genomic location (-a--best--strata). We suppressed reads with more than 50 possible alignments (-m 50). On average, ~97% of reads aligned to the genome (min 90%), of which 59% overlapped a known miRNA. Due to their reduced size, miRNAs are known to be susceptible to cross-mapping, i.e. spurious read alignments to other related miRNAs with strong sequence similarity (de Hoon et al. 2010). In the present dataset, around 65% of reads aligning to known miRNAs had more than one possible alignment on the genome. To mitigate the impact of such cross mapping on miRNA quantification, we used a correction strategy that assigns weights to each of the candidate mapping loci of multiply aligning reads, based on local expression levels and mismatches in the alignment (de Hoon et al. 2010), allowing to distinguish true miRNA reads from likely alignment errors.

### Quantification of miRNA expression

We extracted reads aligning to annotated mature miRNA sequences (miRBase v20) (Kozomara and Griffiths-Jones 2014) with at least 75% overlap using BEDTools (Quinlan and Hall 2010), and divided counts per million associated of each miRNA by the total number of miRNA mapping reads to obtain comparable numbers across all libraries. In addition, we used DESeq2 (version 1.20) (Love et al. 2014) to compute size factors associated to each library and normalize miRNA counts per million across libraries. We then removed lowly, or sporadically, expressed miRNAs by keeping only those with counts of greater than 1 read per million on average across all experimental conditions, leading to a final set of 658 miRNAs across 736 loci. We then added a pseudo-count of 1 RPM to all miRNAs, and log_2_ transformed the data to stabilize the variance of miRNA expression. Linear models were then used to adjust log_2_ transformed counts for technical confounders, such as mean read length of the library (after clipping), or mean GC content of miRNA-aligned reads. Batch effect induced by date of experiment and library preparation were sequentially removed using ComBat (Johnson et al. 2007).

### Assessment of isomiR diversity

For the analyses at the isomiR level, reads aligning to annotated mature miRNA sequences were extracted as described above, and each unique sequence with a mean expression of >1 count per sample was treated as a separate isomiR. MiRNA sequences presenting less than 1 count per sample on average were discarded, and read counts were normalized using the same approach applied for total miRNA expression. We also removed reads where at least one nucleotide could not be called. For each miRNA, the canonical sequence was defined according to miRBase v.20 (Kozomara and Griffiths-Jones 2014) and similarity with canonical sequence at nucleotides 2-7 was used to distinguish *canonical seed* isomiRs from *non-canonical seed* isomiRs. We next classified isoform modifications into three main categories, each subdivided into subtypes of miRNA modifications: (i) *changes in start site*, subdivided in *5’ extension* and *5’ reduction*; (ii) *template changes in end site*, subdivided in *3’ extension* and *3’ reduction*; and (iii) *non-template 3’ additions*, subdivided into *3’ adenylation* and *3’uridylation*. Finally, we quantified, for each miRNA, the frequency of each type of modification, and used these quantities for all downstream analyses. These frequencies were then averaged across miRNAs, to provide global estimates of the frequency of miRNA modification events across samples. After the initial quantification of isomiR diversity, isomiRs that differed only by an internal substitution or a non-canonical terminal addition were merged for downstream analysis, to reduce the effect of sequencing errors.

### Quantification of gene expression levels and transcription rate

RNA-seq reads were aligned to hg19 using Tophat2 (Trapnell et al. 2012) and gene expression values (FPKM) were computed with CuffDiff (Trapnell et al. 2012) based on Ensembl v70. Samples with uneven gene coverage were excluded leaving a total 969 samples with both miRNAs and protein-coding gene expression. Gene expression values were log transformed (with an offset of 1) and corrected for GC content and 5’/3’coverage biais, as well as experiment and library preparation date using linear models and ComBat (Leek et al. 2012). Only the 12,578 genes with mean FPKM>1 where kept for downstream analyses. Further details on gene expression quantification, QC and normalization can be found elsewhere (Quach et al. 2016). Transcription rates were estimated based on the number of nascent unspliced transcripts (Gaidatzis et al. 2015). Namely, for each gene, we used HT-Seq (Anders et al. 2015) to compute the average number of reads mapping to the gene after exclusion of all exonic regions. This number of intronic reads was then divided by the total length of introns, to yield a mean intronic coverage that was used as a proxy of the transcription rate. For each gene, inverse-normal rank transformation was applied to gene expression levels and transcription rate to reduce the impact of outlier values in downstream analyses.

### Differential expression and isomiR analysis

To identify miRNAs that are differentially expressed upon stimulation, we transformed miRNAs counts using an inverse normal rank-transformation, and fitted a linear mixed model of the form 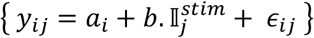, where *y*_*ij*_ is the transformed counts of individual *i* in condition *j, a*_*i*_ is a random effect capturing the inter-individual variability in miRNA expression, *b* is the effect of stimulation on miRNA expression, 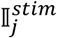 is an indicator variable equal to 0 for the non stimulated samples and 1 for stimulated samples, and *ϵ*_*ij*_ are the residuals. Significance was assessed by maximum likelihood test, and a global Benjamini and Hochberg FDR-correction across all 4 stimuli. Only changes in miRNA expression with corrected p-value < 0.01 were considered significant. To detect significant change in isomiR ratios, we employed a similar approach using isomiRs ratios instead of miRNA read counts.

### Sharing of effects across conditions

To assess the similarity of miRNA response across stimuli, we focused on all miRNAs and isomiRs that were detected to respond to stimulation in at least one condition. We then used a Likelihood-based Model Selection framework (Ding et al. 2018), assuming that miRNAs respond to only a subset of stimuli, and identified the most likely subset of stimuli by jointly modelling rank-transformed miRNA expression, or isomiR ratios, across all 5 conditions. Specifically, for each stimulus j, we assigned an indicator variable *γ*_*j*_ equal to 1 if a given miRNA responds to the stimulus and 0 otherwise. Then, for each of the 15 non-null combinations of stimuli (γ_*j*_)_*j*∈{1,2,3,4}_, we fitted a linear mixed model, as previously performed in each condition {*y*_*ij*_ = *a*_*i*_ + *b. γ*_*j*_ + *ϵ*_*ij*_}, with *a*_*i*_ a random effect capturing the inter-individual variability in miRNA expression or isomiR ratio, *b* is the effect of stimulation and *ϵ*_*ij*_ the residuals, and assigned a probability to each model *m* as

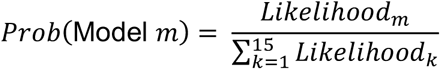

### Detection of miRNA-QTLs and isomiR-QTLs

To identify genetic variants associated with miRNA expression or isomiR ratios, i.e., miR-QTLs and isomiR-QTLs, we focused on the 598 miRNAs that could be uniquely assigned to a single genomic locus and considered a set of 9,854,620 genetic variants with a minor allele frequency (MAF) > 0.05 in either African- and European-ancestry groups, of which 1,981,401 were located < 1Mb from one the 598 mature miRNAs. We used *MatrixEQTL* (Shabalin 2012) to map miR-QTLs within a 1Mb window on each side of mature miRNAs. miR-QTL mapping was performed separately for each condition, merging both populations and including an indicator variable to control for the effect of population on miRNA expression. MiRNA counts per million values and isomiR ratios were rank-transformed to a normal distribution before mapping, to reduce the impact of outliers. FDR was computed by mapping miR/isomiR-QTLs on 100 permuted datasets, in which genotypes were randomly permuted within each population. We then kept, for each permutated dataset, the most significant *p*-value per miRNA or isomiR, across all conditions, and computed the FDR associated with various *p*-value thresholds ranging from 10^−3^ to 10^−50^. We subsequently selected the *p*-value threshold that provided a 5% FDR (*p* <10^−6^).

### Comparison of miR-QTLs to protein-coding gene eQTLs

To compare the degree of genetic control of miRNA expression variability to that of protein-coding genes and long non-coding RNAs, we used the *MatrixEQTL* package (Shabalin 2012) to perform eQTL analyses, as performed for miR-QTLs. We considered a 1Mb window around the TSS of each gene, and tested all frequent SNPs (i.e. with MAF>5% in either population) for association with gene expression. eQTLs were tested in both populations combined on rank transformed gene expression values, adjusting for population. FDR was computed based on 100 permutations as described for miRNAs. When comparing frequency of eQTLs and miR-QTLs, a joint FDR was re-computed considering miRNAs, protein-coding genes and long noncoding RNAs together, to ensure similar power for the detection of miR-QTLs and eQTLs. Protein-coding genes were then assigned to various categories based on Exac pLI scores (LOF intolerant – pLI > 0. 9, Recessive pRec>.9, neutral pNull>.9) (Lek et al. 2016) or GO annotations (TF – GO:0003700). In addition, to ensure that differences in read coverage between miRNAs and protein-coding genes were not responsible for a lower power to detect miR-QTLs/eQTLs, we evaluated the impact of read coverage on the detection of miR-QTL and eQTLs (Supplemental Fig. S4A,B). We removed from our analyses miRNAs and genes with < 50 supporting reads on average.

### Sharing of miR-QTLs and eQTLs across conditions

When comparing miR-QTLs across conditions, we used a likelihood-based model selection framework to increase power for detection of shared effects. Namely, for each SNP-miRNA pair, rank-transformed miRNA expression, or isomiR ratios, *y*_*ij*_ were modelled jointly across all 5 conditions. An indicator variable *γ*_*j*_ was defined as 1 if the miRNA is under genetic control in condition j and 0 otherwise. Then, for each of the 31 non-null combinations of stimuli (γ_*j*_)_*j*∈{1,2,3,4,5}_, we fitted a linear model of the form {*y*_*ij*_ = *a*_*jp*_ + *b. γ*_*j*_*SNP*_*i*_ + *ϵ*_*ij*_}, with *a*_*jp*_ the mean expression of the miRNA or isomiR in condition *j* and population *p*, SNP_i_ the number of minor alleles carried by individual *i, b* the mean effect of the SNP in conditions where it is active and *ϵ*_*ij*_ the residuals. Each model was then assigned a probability as: 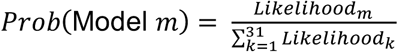, and the model with the highest probability was retained. The same approach was applied to assess the sharing of eQTLs across conditions, using rank transformed mRNA levels (FPKM) of protein-coding genes instead of miRNA expression. In addition, to identify response-miR-QTLs, we tested for significant differences in effect size of miR-QTLs between the stimulated and non-stimulated state, using an interaction test. Rank-transformed miRNA expression, or isomiR ratios, *y*_*ij*_ are decomposed between *a*_*jp*_ the mean expression of the miRNA or isomiR in condition *j* and population *p*, the effect of the SNP at basal state *b*, and the differences in effect size between basal and stimulated state *c.*

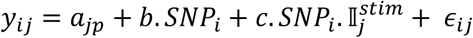

The significance of the interaction was then tested by a Student t test for H_0_: {*c=0*}.

### Annotation of miRNA TSS and miR-QTLs

Transcription start site (TSS) of miRNAs were obtained from Fantom5 data, based on (de Rie et al. 2017), together with their conservation levels (mean PhastCons of promoter region). Hairpin coordinates were retrieved from mirBase V20 (Kozomara and Griffiths-Jones 2014). MiR-QTLs for which TSS information was available were then classified based on their location relative to the TSS and hairpin. Namely, miR-QTLs were first classified as *miRNA-altering* or *hairpin-altering* if they overlapped the sequence of the mature miRNA or its associated hairpin. Then, we computed, for each miR-QTL, the distance between the SNP and both the hairpin and the TSS of the associated pri-miRNA. miR-QTLs that were located less than 20kb from the TSS or the hairpin were annotated as *hairpin-* or *TSS-flanking*, according to the feature from which they were the closest. Finally, miR-QTLs located > 20kb from both TSS and hairpin, were annotated as *Distant*. Overlap of miR-QTLs (and eQTLs) with promoters and enhancers was assessed based on Epigenomic Roadmap data, using the ChromHMM segmentation of tissue E029 (CD14^+^ monocytes) (Roadmap Epigenomics Consortium, 2015). Specifically, regions assigned to class 1 and 2 (Active TSS and Active TSS Flank) were considered as promoters, and class 6 and 7 (Enhancers and Genic enhancers) were considered as enhancers. miR-QTL and eQTL peak SNPs were then compared to the set of all frequent SNPs (MAF>5%) considered in our study.

### Population differences in miRNA and isomiR expression

To identify miRNAs that are differentially expressed between populations, we applied Student’s t-test to inverse normal rank-transformed miRNAs counts within each condition separately, comparing African-to European-ancestry individuals. A global Benjamini and Hochberg FDR-correction was applied across all 5 conditions to evaluate significance. Only changes in miRNA expression with corrected p-value < 0.01 were considered as significant. A similar approach was used to test for population differences in isomiR levels, using isomiRs ratios, instead of miRNA read counts. Sharing of population differences among conditions was assessed using a model selection framework similar to the one used to assess sharing of mir-QTLs. For each individual *i* and condition *j*, we assigned an indicator variable *γ*_*j*_ equal to 1 if a miRNA is differentially expressed between populations in that condition and 0 otherwise. Then, for each of the 31 non-null combinations of conditions (γ_*j*_)_*j*∈{1,2,3,4,5}_, we fitted a linear model 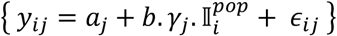, with *a*_*j*_ a the mean expression of the miRNA in condition *j* across African-ancestry individuals, *b* the mean difference in miRNA expression between European- and African-ancestry individuals, and *ϵ*_*ij*_ a normally distributed residual. Each possible model is then assigned a probability *m* as 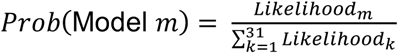 and the most likely model is retained.

### Assignment of miRNA targets

miRNA targets were predicted using miRanda v3.3a (Enright et al. 2003), providing canonical sequences obtained from miRBase V20 (Kozomara and Griffiths-Jones 2014) as input and 3’UTR sequence of known transcripts based on Ensembl V70. Defaults settings were used for target prediction, and a gene was considered as targeted by a miRNA if at least one of its annotated transcripts had a predicted binding site for the miRNA. Prediction of isomiR targets was performed in a similar manner, using the isomiR sequence instead of the canonical sequence.

### Assessment of miRNA-mRNA correlations

To identify likely miRNA-gene interactions occurring in each condition, we modelled gene expression as a function of miRNA levels, using population as a covariate. All miRNAs were introduced simultaneously in the model and an elastic net penalty (Friedman et al. 2010) was set on the miRNA effects to make the model identifiable, leading to the following model

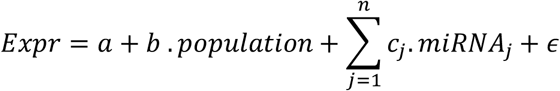

with 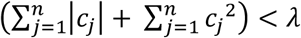.

Here, *Expr* is the vector of gene expression across all samples from the condition under study, *population* is an indicator variable representing population of origin, (*miRNA*_*j*_)_*j=1..n*_ are the vectors of expression of the 658 expressed miRNAs, and *ϵ* is a random Gaussian noise. *a* denotes the mean expression in the reference population, *b* and (*c*_*j*_)_*j=1..n*_, are parameters capturing the effect of population, and miRNAs on gene expression. *λ* is a constant value that captures the amount on constraint on the effect of miRNAs included in the model.

Using this model, we performed stability selection (Hofner et al. 2015) to select miRNAs that have a significant effect on gene expression with high probability. Briefly, stability selection consists in performing repeated sub-samplings of the data, typically considering only half of the initial data, and selecting the first Q miRNAs with non-null c_j_ coefficients across increasing values of *λ*. Under the reasoning that only miRNAs with a true effect on gene expression will be consistently selected across subsamplings, we can then use the frequency at which a miRNA is kept in the model as the posterior probability that this miRNA has a significant impact on gene expression. We performed 100 resamplings with varying values of Q (from 3 to 60, by steps of 3) and estimated, for each value of Q, the probability of each miRNA to be included in the model. Then, for each value of Q, we randomly permuted the data and repeated the procedure to obtain the distribution of inclusion probabilities, under a model where gene expression is independent from miRNA levels. A single permutation of the data was performed for each Q, and the null distribution was estimated across all 12,578 genes × 658 miRNAs. Based on this null distribution, we computed the FDR associated with a given probability threshold, as the ratio between the number of significant gene-miRNA pairs that exceed that probability threshold in the permuted and non-permuted data set. We found that setting Q=9 maximized the number of significant associations at an FDR of either 1%, 5% of 10%, and used this value for all subsequent analyses. We then considered as significant all miRNAs that reach a posterior probability of 0.8, which is equivalent to an FDR of ~1.3% based on our permutation setting (Supplemental Fig. S5).

### Correlation between miRNAs and transcription

Correlation between miRNAs and transcription rate was obtained using the *MatrixEQTL* package (Shabalin 2012), providing miRNAs instead of genotypes and adjusting for population. All associations where the miRNA was located less than 1kb away from the gene were discarded, and a 5% FDR was used to declare associations as significant. For each miRNA, significantly associated genes were split into *negatively* and *positively* correlated genes according to the sign of the corresponding β parameter. We then tested each set of associated genes for enrichment in predicted binding sites obtained from miRanda (Enright et al. 2003) compared to the set of all transcribed genes with at least one predicted miRNA binding site. Benjamini-Hochberg correction was applied across all miRNAs for both positive and negative correlations, and only enrichments passing a 5% FDR were retained.

### *Trans*-effects of miRNA-QTLs on gene expression

To assess the effect of miR-QTLs and isomiR-QTLs on gene expression, we considered the set of 118 unique SNPs with an effect on miRNA expression or isomiRs ratios, and tested for *trans*-associations with genes located >1Mb away from these SNPs. We then used the pi0est function from the qvalue package to estimate the percentage π_1_^obs^ of genes associated with a miR-QTL or isomiR-QTL across all 5 conditions, based on the shape of the *p*-value distribution. Finally, we repeated the same analysis for 100 random samples of SNPs, matched for minor allele frequency (using MAF bins of 5%) and computed a resampling *p*-value by counting the frequency at which the percentage π_1_^obs^ of genes associated to miR-QTLs and isomiR-QTLs exceeded the π_1_ value estimated from sets of randomly selected SNPs.

### Relative contribution of transcription and miRNAs to gene expression variability

To account for co-transcription when assessing miRNA-gene correlations, we repeated our stability selection approach adding transcription as a covariate in the model. The final model can thus be written as

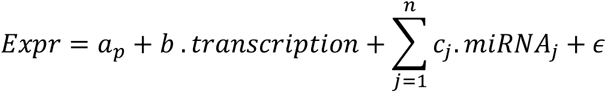

With 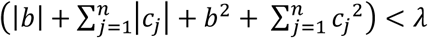.

Here, *Expr* and *transcription* are the vectors of gene expression and transcription rate across all samples from the condition under study, (*miRNA*_*j*_)_*j=1..n*_ are the vectors of expression of the 658 expressed miRNAs, and *ϵ* is a random Gaussian noise. *a*_*p*_ denotes the mean expression in population *p* and *b* and (*c*_*j*_,)_j=1..n_ are parameters capturing the effect of transcription and miRNAs on gene expression. *λ* is a constant value that captures the amount on constraint on miRNAs that are included in the model.

After identifying miRNAs that have a significant effect on gene expression, miRNA effect sizes were assessed using CAR scores as implemented in the care package (Zuber and Strimmer 2011). Briefly, CAR scores (noted *ω*) are a variation of partial correlations that allows to measure correlations between one or more covariates and a response variable, while adjusting each covariate for the effect all other covariates. More importantly, the squared CAR scores (*ω*^2^) sum to the total percentage of variance explained by the model (R^2^), allowing to interpret the square of each individual CAR score as the percentage of variance explained by the associated covariate, when adjusting for all other covariates. To evaluate the variance explained by a subset of miRNAs (i.e. negatively correlated miRNAs or negatively correlated miRNAs with a known binding site to the gene), we considered the sum of *ω*^2^ over all miRNAs of that subset (using the sign of *ω*, to identify negative correlations).

## DATA ACCESS

The miRNA-sequencing data generated in this study have been deposited in the European Genome-phenome Archive (EGA) under accession code EGAS00001004192.

## ACKNOWLEDGEMENTS

The laboratory of L.Q.-M. is supported by the Institut Pasteur, the Collège de France, the French Government’s *Investissement d’Avenir* program, *Laboratoires d’Excellence* “Integrative Biology of Emerging Infectious Diseases” (ANR-10-LABX-62-IBEID) and “Milieu Intérieur” (ANR-10-LABX-69-01), and the *Fondation pour la Recherche Médicale* (Equipe FRM DEQ20180339214). This project was funded by the European Research Council under the European Union’s Seventh Framework Programme (FP/2007–2013)/ERC grant agreement 281297. We thank Macrogen Inc. for the use of their RNA-sequencing facilities.

## Author contributions

M.R. and K.J.S conceived the analysis pipeline, M.R. supervised the analyses, M.R., M.S. and K.J.S. analyzed and interpreted the data, J.P. and H.Q. designed and performed the experiments; L.Q.M. interpreted the data, conceived and supervised the study, and obtained funding, M.R. and L.Q.M. wrote the manuscript, with inputs from all authors. All authors read and approved the final manuscript.

## Supplemental Figures

**Supplemental Figure S1**. Quality and pre-processing of small RNA sequencing data.

**Supplemental Figure S2**. IsomiR diversity and rate of miRNA modifications.

**Supplemental Figure S3**. Sources of isomiR variation upon immune activation.

**Supplemental Figure S4**. Genetic basis of miRNA expression.

**Supplemental Figure S5**. Detection of miRNA-mRNA correlations.

## List of Supplemental Tables

**Supplemental Table S1**. (*A*) List of all 736 loci encoding a miRNA, with their coordinates and the name of the associated miRNA. Coordinates and ID of the associated hairpin and transcription start site (TSS) are provided. (*B*) List of all 2,359 frequent isomiRs sequences with their associated miRNA and hairpin. Details of observed deviations from the canonical isomiR are provided. (*C*) Number of isomiRs per miRNA at a frequency above 1% or 5%, before and after merging isomiRs that differ only by non-terminal substitutions. (*D*) For each type of miRNA modification, average frequency of occurrence of the modification among 3p and 5p miRNAs and significance of the differences in frequency between both arms (Wilcoxon test).

**Supplemental Table S2**. (*A*) List of 340 miRNAs that change their expression in response to at least one stimulus. For each miRNA, non-stimulated expression and fold changes upon stimulation are provided, together with the significance of expression changes and the inferred model of sharing across conditions. (*B*) List of isomiRs that change their ratios upon stimulation. For each isomiR, the name of the associated miRNA is reported together with the % of miRNA reads that it accounts for in the non-stimulated state. For each stimulus, changes in expression ratio are indicated together with their associated p-value. The inferred model of sharing across conditions is provided. (*C*) For each type of miRNA modification, changes in frequency of occurrence of the modification upon stimulation, and their significance. Average frequency in the non-stimulated state is also reported. Results are provided for all miRNAs together (any) or focusing on 3p or 5p arm miRNAs.

**Supplemental Table S3**. (*A*) List of 122 miRNAs associated with at least one miR-QTL, and their most strongly associated variant. Association statistics (β coefficient, *p*-value, R^2^) are provided for each condition. For each SNP, genomic coordinates are reported, together with derived allele frequency in African- and European-ancestry individuals. miR-QTL classification based on the distance from the mature miRNA, pre-miRNA and TSS (when available) is also provided (miRNA-altering, hairpin-altering, hairpin-flanking, TSS-flanking, and Distant) as well as overlap with CD14^+^ promoters/enhancers based on Epigenomic Roadmap. (*B*) Tests for genotype×stimulation interactions across all 4 stimuli for the 122 miR-QTLs. Association statistics (interaction coefficient, *p*-value) are provided for each condition, and the best model of sharing of the miR-QTL is reported. (*C*) List of 25 isomiRs associated with at least one isomiR-QTL, and their most strongly associated SNP. For each SNP, genomic coordinates are reported, together with the derived allele frequency (DAF) in African- and European-ancestry groups.

**Supplemental Table S4**. (*A*) List of 244 miRNAs that are differentially expressed between populations in at least one condition. For each miRNA and condition, average expression in African- and European-ancestry groups is provided, together with significance of differences in expression. The inferred model of sharing of these differences across conditions is provided. (*B*) List of 188 isomiRs that are differentially expressed between populations. For each isomiR and condition, average ratio in African- and European-ancestry groups is provided, together with significance of differences between populations. The inferred model of sharing of these differences across conditions is also provided. (*C*) Tests for population×stimulation interactions across all 4 stimuli for all 658 tested miRNAs. Association statistics (mean logFC in each population and interaction *p*-value) are provided for each stimulus. The best model of sharing of the popDE is reported for the 244 miRNAs that are differentially expressed between populations in at least one condition. (*D*) For each type of miRNA modification and conditions, differences in frequency of occurrence of the modification between populations and their significance. Results are provided for all miRNAs together (any) or focusing on 3p or 5p arm miRNAs.

